# Parabrachial bombesin receptor subtype 3 neurons facilitate heat hypersensitivity in persistent pain

**DOI:** 10.1101/2025.03.05.641630

**Authors:** Heather N. Allen, Erick J. Rodríguez-Palma, Tyler S. Nelson, Naomi K. Grabus, Nia A. Dufeal, Rajesh Khanna

**Affiliations:** Department of Pharmacology & Therapeutics, McKnight Brain Institute, Center for Advanced Pain Therapeutics and Research (CAPToR), College of Medicine, University of Florida, Gainesville, FL, 32610, USA

**Keywords:** pain, inflammation, neuropathic, parabrachial nucleus, bombesin receptor subtype 3

## Abstract

The parabrachial nucleus (PBN) is a critical hub for pain processing that acts as a switchboard for nociceptive signals, relaying sensory information to forebrain regions that integrate the sensory and affective dimensions of pain. Although the PBN is well established as a key regulator of pain, the remarkable heterogeneity of its neuronal populations has hindered efforts to identify specific cell types responsible for distinct aspects of pain processing. Here, we identify bombesin receptor subtype 3 (*Brs3*)-expressing neurons as a distinct glutamatergic PBN subpopulation involved in heat hypersensitivity associated with persistent pain. Using *Fos* expression analysis and *in vivo* calcium imaging, we demonstrate that *Brs3* neurons exhibit heightened activity in response to heat stimulation following an inflammatory insult or neuropathic injury. Inhibition of *Brs3* neurons effectively reduces heat, but not mechanical, hypersensitivity induced by both inflammatory and neuropathic pain, suggesting a specific role in processing heat hypersensitivity. Ablation of parabrachial *Brs3* neurons prior to induction of pain also selectively prevents the development of heat hypersensitivity induced by persistent inflammation in mice. *Brs3*-expressing neurons encompass multiple previously identified pain-related PBN subpopulations, including those expressing the mu opioid receptor (*Oprm1*), tachykinin 1 receptor (*Tacr1*), and neuropeptide Y Y1 receptor (*Npy1r*), positioning *Brs3* as a potential unifying marker of heat hypersensitivity circuits. These findings provide new insight into the organization of pain-processing networks in the PBN and highlight *Brs3* neurons as a crucial population for heat pain.

## Introduction

Pain is a multifaceted experience and essential protective mechanism that helps the body maintain homeostasis and avoid injury^1^. The International Association for the Study of Pain’s most recent definition of pain emphasizes that “pain cannot be inferred solely from activity in sensory neurons”, acknowledging the necessity of supraspinal circuits in creating and shaping the aversive nature of the pain experience^2^. Although pain often originates in the periphery, its perception depends on intricate supraspinal processing, where emotional and cognitive inputs are integrated with physical nociceptive signals to ultimately produce the unpleasant sensation of pain^3^.

Persistent pain is driven by a variety of disease processes that induce lasting changes in peripheral and central nociceptive pathways^4^. Inflammatory pain, the result of tissue injury and immune activation, is a hallmark of many chronic pain conditions^5^, including arthritis^6^, post-surgical pain^7^, low back pain^8^, fibromyalgia^9^, nerve damage^10^, endometriosis^11^, and inflammatory bowel disease^12^. Persistent inflammatory pain can lead to prolonged hypersensitivity and maladaptive changes to nociceptive circuits, contributing to the transition from acute to chronic pain^13,14^. Similarly, neuropathic pain arises from damage or disease to the nerves and results in hyperexcitability of the sensory neurons in the dorsal root ganglia, loss of spinal inhibition, and activation of central sensitization pathways^15–18^. Although peripheral in origin, neuropathic pain drives reorganizational changes in supraspinal circuits as it persists^4,19,20^. A deeper understanding of how enduring pain alters pain circuits is critical for designing effective therapeutics.

The parabrachial nucleus (PBN) is a major hindbrain hub for pain processing, relaying ascending nociceptive information from via the spinal cord to forebrain regions involved in the sensory and affective dimensions of pain^21^. In chronic pain conditions, PBN neurons exhibit heightened activity in response to noxious stimuli, contributing to the amplification of pain observed in both neuropathic^22–27^ and inflammatory^27–29^ pain conditions. Glutamatergic neurons, which comprise approximately 90% of all PBN neurons^30^, are particularly important contributors to pain modulation: they are activated by inflammatory and neuropathic pain conditions^23,31–33^, and their activation alone is sufficient to induce hypersensitivity in mice, even in the absence of injury^23^. Despite their central role in pain, glutamatergic PBN neurons comprise dozens of molecularly distinct populations, making it difficult to identify the circuit architecture underlying pain modulation^30^. Several distinct glutamatergic subpopulations have been directly implicated in pain processing. PBN neurons expressing the mu opioid receptor^34^, tachykinin 1 receptor^35^, or neuropeptide Y Y1 receptor-expressing neurons^36–38^ have each separately been shown to contribute to the modulation of inflammatory and/or neuropathic pain in mice. However, while some co-expression exists between these populations^30,36,39^, they exhibit significant anatomical and molecular divergence, making it challenging to determine how they synchronize to modulate pain in the PBN.

Recent spatial transcriptomic analyses suggest that one molecular population may unify several previously identified pain-modulating PBN neuron types: neurons expressing bombesin receptor subtype 3 (*Brs3*)^30^. Brs3, is an orphan Gq-coupled receptor with no identified endogenous ligand, and its physiological function remains poorly understood^40^. However, *Brs3* expression in the PBN marks a distinct subset of neurons that encompasses the separate mu opioid receptor-, tachykinin 1 receptor-, and neuropeptide Y Y1 receptor-expressing PBN populations^30^. These findings raise the possibility that *Brs3* neurons function as a common integrative node coordinating multiple pain-modulating PBN populations. Here, we tested whether *Brs3*-expressing PBN neurons regulate persistent pain using complementary molecular, genetic, and behavioral approaches in mouse models of inflammatory and neuropathic pain.

## Methods

### Animals

Male and female C57Bl/6J (Jackson labs, #000664) and *Brs3^Cre^* mice (B6.Cg-*Brs3^tm^*^1*.1(cre/GFP)Rpa/J*^; Jackson labs, #030540)^41^ aged 6-10 weeks were used for all experiments. Animals were group housed in a temperature and humidity-controlled room on a 12:12 hour light:dark cycle with *ad libitum* access to food and water. All procedures were approved by the Institutional Animal Care and Use Committee at the University of Florida (IACUC protocol #202400000002). All experiments included both male and female animals.

### Surgery

Male and female *Brs3^Cre^* (*Brs3^Cre^*, *Brs3^Cre/+^* and *Brs3^Cre/Cre^*) and *Brs3^+/+^* wildtype mice were anesthetized using isoflurane (5% induction, 2.5% maintenance) and had their heads fixed in a stereotax and a small incision made in the scalp. For fiber photometry experiments, a craniotomy was performed over the right PBN (coordinates A/P -5.15 mm, M/L +/-1.1 mm, D/V -3.35 mm) and 300 nL of a Cre-dependent virus carrying a genetically encoded calcium indicator (AAV9-hSyn-flex-GCamp6s, #100845, Addgene, titer 1.8 x 10^13^ GC/mL, Watertown, Massachusetts, USA) was injected at a rate of 1 nL/sec using a Nanoject III programmable nanoliter injector (Drummond Scientific, Broomall, PA, USA) connected to a glass pipette. The injector was left in place for an additional five minutes to prevent backflow and to allow the virus to diffuse. For fiberoptic implant surgeries, one bone screw was attached to the skull left of midline anterior to lambda and one bone screw was attached right of midline anterior to bregma. A fiberoptic implant (RWD Life Science, San Diego, CA, USA; 1.25 mm ferrule diameter, 200 mm core, 0.37 NA) was implanted above the injection site in the right PBN and secured to the skull using dental cement. For chemogenetic experiments, bilateral craniotomies were performed over the left and right PBN and 300 nL of Cre-dependent virus carrying an inhibitory (AAV8-hSyn-DIO-hM4Di-mCherry, titer 2.8 x 10^13^ GC/mL, #44362 Addgene) construct was injected at a rate of 1 nL/s into *Brs3^Cre^* and *Brs3*^+/+^ animals. For genetic ablation experiments, 300nL of a virus carrying Cre-dependent diphtheria toxin A (AAV2/8-Ef1a-mCherry-flex-dtA, titer 5.4 x 10^12^ GC/mL, COVF Viral Vector, Québec City, QC, Canada) was injected into the left and right PBN of *Brs3^Cre^* and *Brs3*^+/+^ animals at a rate of 1 nL/sec. The injector was left in place for an additional five minutes to prevent backflow and to allow the virus to diffuse. The scalp was closed using 6.0 silk suture and triple antibiotic ointment was applied to the wound. Animals received a subcutaneous injection of Meloxicam (20 mg/kg, Patterson Veterinary Supply, Alachua, FL, USA) and recovered on a heating pad before being returned to their homecage, where they received acetaminophen water (1.1 mg/mL) for 72 hours post-surgery^42^. Mice were allowed at least three weeks of recovery post brain surgery before undergoing behavior experiments.

Spinal nerve ligation was performed as previously described^24^. Briefly, mice were anesthetized with isoflurane (5% induction, 2.5% maintenance), and the paraspinal muscles were separated at the L4 level and the L5 transverse process was carefully removed. The left L4 and L5 spinal nerves were exposed and tightly ligated with 6-0 silk suture distal to the dorsal root ganglion. Sham animals underwent an identical surgery where the spinal nerves were exposed but not ligated. Animals recovered on a heating pad after surgery.

### Drugs

Under 2% isoflurane anesthesia, 20 µL of undiluted Complete Freund’s Adjuvant (CFA, #AR001, Millipore Sigma, Burlington, MA, USA,) was injected into the plantar surface of the left hindpaw using a 30 gauge needle attached to a 25 µL Hamilton syringe. Control animals were anesthetized with 2% isoflurane but received no injection in the hindpaw.

For chemogenetics, compound 21: 11-(1-Piperazinyl)-5H-dibenzo[b,e][1,4]diazepine dihydrochloride, also called DREADD agonist 21 dihydrochloride (C21), CAS 2250025-92-2(#HB6124, Hello Bio, Princeton, NJ, USA), was diluted in saline the day of the experiment and injected intraperitoneally (1 mg/kg) 30 minutes before behavioral testing.

### Behavior

All behavior was performed blinded to drug injection (C21 or saline) and/or virus (hM4Di, hM3Dq, mCherry). All experiments included both male and female mice.

*Fiber photometry*. Fiber photometry experiments were conducted as previously described^24,25,36,43,44^. Animals were habituated to Plexiglas chambers, elevated wire mesh, and patch cable attachment for 1 hour each two days prior to experimentation. On the day of experimentation, animals were habituated in Plexiglas chambers on elevated wire mesh for one hour before having patch cables attached to the implanted fiberoptics and habituating for an additional 30 minutes with the experimenter in the room. Baseline recordings were performed to innocuous and noxious stimuli according to the following paradigm: 0.07 g von Frey filament, 1.0 g von Frey filament, radiant heat, and blunted pin prick. Calcium transients were collected continuously (FP3002, Neurophotometrics, San Diego, CA, USA) during the stimulation protocol. Each stimulus was applied to the plantar surface of the left hindpaw three times two minutes apart, and all three responses were averaged to represent the animal’s response to that stimulus. Custom MatLab scripts were used to normalize the GCamp6s (470 nm) signal to the isobestic signal (405 nm), controlling for motion artifacts as well as photobleaching. The change in fluorescence (dF/F) was calculated by subtracting the GCamp6s signal during stimulation from the average GCamp6s signal over the 15 seconds directly prior to stimulation. The area under the curve (AUC) was calculated for the five seconds directly following stimulus application. Following baseline recordings, animals received a CFA injection in the left hindpaw or left leg SNL, and the fiber photometry protocol was repeated 3 days after CFA or 21 days after SNL. All animals were perfused following the experiment for histological verification of viral injection and fiber optic placement in the PBN.

*Static mechanical sensitivity*. Mechanical withdrawal thresholds were determined using the von Frey assay as previously described^45^. Mice were habituated to Plexiglas chambers on an elevated wire mesh platform for 1.5 hours prior to testing. Calibrated von Frey filaments (#58011, Braintree Scientific, Braintree, MA, USA) were used to assess mechanical sensitivity at the plantar surface of the left hindpaw using the up/down method^46^.

*Dynamic mechanical sensitivity*. Following von Frey testing, dynamic mechanical sensitivity was tested while mice were in the same chambers. A cotton swab was teased apart so that the cotton was approximately three times its original size, and the hindpaw was lightly stroked from heel to toe. The stimulation was repeated four times, and the percent of withdrawals was reported as response frequency^37,47^.

*Cold withdrawal latency.* Directly following mechanical sensitivity testing, mice were assessed for cold sensitivity using the acetone droplet test^45^. Using a 5 mL syringe attached to PE-90 tubing with a flared end, approximately 10 µL of acetone was applied to the plantar surface of the left hindpaw. The time in seconds the animal spent licking, biting, and attending to the paw was measured. The test was repeated three times, and the average of all three trials is reported.

*Pinprick assay*. Following cold withdrawal latency, animals were tested in the pinprick assay. A 27-gauge needle was applied to the plantar surface of the hindpaw without piercing the skin. Paw withdrawal, shaking, and licking were considered a positive response. The stimulation was repeated four times, and the percent of withdrawals was reported as response frequency^47^.

*Heat withdrawal latency.* Thermal withdrawal latency was assessed using the hotplate^36^ and the Hargreaves Apparatus^48^. For hotplate, mice were gently placed on a 52°C Ugo Basile hot/cold plate within an acrylic enclosure (#55075, Stoelting, Wood Dale, IL, USA). The time until heat withdrawal response, defined as licking, jumping, or flinching, was recorded in seconds. The animal was immediately removed following withdrawal or after a cutoff of 30 seconds to avoid tissue damage. Three trials were averaged with a between-trial interval of at least ten minutes. For Hargreaves, mice were habituated in Plexiglas testing chambers placed on an elevated glass platform of the Ugo Basile Thermal Plantar apparatus (#55370, Stoeling, Wood Dale, IL, USA) for 1 hour. Radiant heat was applied to the center plantar surface of the animal’s left paw through the glass bottom. The time until the hindpaw withdrawal was recorded in seconds, with a cutoff of 15 seconds. Three trials were averaged with a between-trial interval of at least 10 minutes.

### Fluorescence *in situ* hybridization

*Stimulus-evoked Fos expression*: Naïve and CFA treated animals were lightly anesthetized and received a heat stimulus to induce *Fos* expression. Half of each group (naïve and CFA) received a heat stimulus, and half received no stimulus (control). Control animals were lightly anesthetized (∼1% isoflurane) for five minutes. Heat stimulated animals were lightly anesthetized (∼1% isoflurane) and had the left paw dipped in 55°C water for ten seconds. Heat stimulation was repeated four times, with a minute between each stimulation. Animals were returned to their homecage for 1 hour before being perfused with ice cold PBS followed by 10% neutral buffered formalin. Tissue was extracted and prepared for *in situ* processing. RNAscope multiplex fluorescent V2 assay was used with probes for *Brs3* (ACD Bio #454111-C3; Advanced Cell Diagnostics (ACD Bio), Newark, CA, USA) or *Fos* (ACD Bio #316921). Brains were extracted and post-fixed in 10% neutral buffered formalin for 2 hours at 4°C before being transferred to 30% sucrose and stored at 4°C for 3 days. Tissue was embedded in optimal cutting temperature gel (OCT, TissueTek) before being sectioned at 20 µm on a cryostat. Three to five representative PBN sections from each animal were mounted on SuperFrost Plus microscope slides (Fisher Scientific) and allowed to airdry overnight at room temperature. Slides were rinsed in MilliQ water for five minutes to remove OCT followed by two consecutive baths of 100% ethanol (2 minutes each). Tissue was treated with Protease III for 20 minutes in HybEZ oven at 40°C. RNAscope was performed according to manufacturer’s instructions (ACD Bio) as previously described^49^. Briefly, tissue was incubated with probes for *Brs3* (ACD Bio #454111-C3) and *Fos* (ACD Bio #316921) for two hours at 40°C before undergoing a series of amplification steps according to manufacturer’s instructions (AMP1 30 min, AMP2 30 min, AMP 3 15 min) at 40°C. Each probe had a TSA-based fluorescent label developed in succession followed by application of DAPI for 30 seconds. Slides were coverslipped with VECTASHIELD hardset anti-fade mounting medium with DAPI (#H-1400-10, Vector Laboratories, Newark, CA, USA) and imaged on a Leica DMI8 microscope (Wetzlar, Germany) using a 20x objective lens. Images were analyzed using QuPath software v0.4.3. The lateral PBN was outlined according to the mouse reference brain atlas (Allen Institute) and cells with greater than three puncta (*Fos, Brs3*) surrounding a DAPI labeled nucleus were considered positive. Cell were considered positive for co-expression if they had greater than three puncta for each marker. Between 3 and 5 sections from across the rostral-caudal axis of the PBN were quantified and averaged for each animal.

*Brs3 ablation confirmation*: Following behavior in dtA injected mice, animals were perfused with ice cold PBS followed by 10% neutral buffered formalin. Tissue was extracted and prepared for *in situ* processing and RNAscope multiplex fluorescent V2 assay for *Brs3* (ACD Bio #454111-C3) was conducted as described above. Cells with greater than three puncta *(Brs3*) surrounding a DAPI labeled nucleus were considered positive. Between 3 and 5 sections from across the rostral-caudal axis of the PBN were stained for each animal.

### Immunohistochemistry

Animals receiving viral injections were transcardially perfused following behavioral experiments with ice cold PBS followed by ice cold 10% neutral buffered formalin. Brains were extracted and post fixed in 10% neutral buffered formalin overnight at 4°C. Brains were then transferred to 30% sucrose at 4°C for five days before being frozen and sectioned at 35 µm on a cryostat. Floating PBN sections were washed 3 times for 5 minutes each in PBS before being incubated in blocking buffer (5% normal goat serum, 0.1% Triton X-100 in PBS) for sixty minutes. Sections were then incubated in a 1:1000 dilution of primary antibody against either GFP (Rb anti-GFP, Millipore Sigma, #AB3080) for fiber photometry animals or mCherry (#ab167453, rabbit anti-mCherry, Abcam, Waltham, MA, USA,) for chemogenetic and dtA animals overnight. Following three 5-minute washes, sections were incubated in secondary antibody goat anti rabbit Alexa Fluor 488 (1:1000 #A11008, Invitrogen, Waltham, MA, USA), or goat anti rabbit Alexa Fluor 594 (1:1000 #A21442, Invitrogen, Waltham, MA, USA) for 90 minutes. After three 5-minute washes in 1% phosphate buffer, sections were mounted on SuperFrost Plus microscope slides and coverslipped with VECTASHIELD hardset mounting medium with DAPI (Vector Laboratories, #H-1400-10). Images were captured on Echo Revolution microscope using a 20x objective lens. Viral infection was confirmed by comparing fluorescently labeled cells to the Allen Brain atlas to align with the PBN.

### Statistics and data analysis

Data were analyzed using GraphPad Prism v10.4.1 and all values are presented as mean ± SEM. Statistical significance is determined as P<0.05. *In situ* data were analyzed using one-way ANOVA and mixed effects analysis with Tukey’s test for multiple comparisons. Fiber photometry data were analyzed using two-tailed paired Student’s T-tests. Inhibitory DREADD and dtA hotplate data were analyzed using two-way analysis of variance (ANOVA) followed by Uncorrected Fisher’s LSD for multiple comparisons. All results were preliminarily tested for sex differences. Data is presented with males and females pooled, as no sex differences were identified.

## Results

Previous studies have demonstrated that neurons in the PBN become sensitized during both inflammatory and neuropathic pain, exhibiting enhanced responses to noxious stimuli^23,25–29,31,48,50–52^. To determine whether *Brs3*-expressing neurons are similarly recruited during persistent inflammatory pain, we used multiplex *in situ* hybridization to quantify co-expression of *Brs3* and *Fos* mRNA in the PBN of naïve and CFA-treated mice following exposure to light anesthesia alone or light anesthesia combined with a noxious heat stimulus applied to the left hindpaw (**Figure 1A**). CFA-treated mice exposed to a heat exhibited displayed a larger percent of *Fos* positive *Brs3* cells in the PBN (38.5%) compared to non-heat stimulated CFA animals (22.4%) and naïve animals exposed to heat (11.6%) or not (6.6%) (**Figure 1B-G**). These findings indicate the *Brs3*-expressing PBN neurons are selectively recruited by noxious heat following inflammatory injury but not under basal conditions.

**Figure 1:**
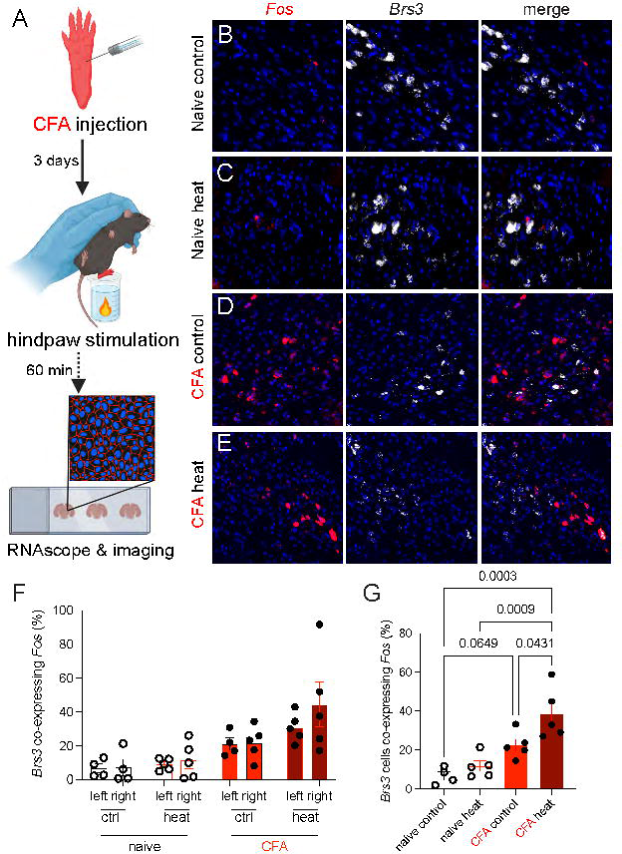
Heat stimulation increases *Fos* expression in parabrachial *Brs3* neurons from CFA animals. (A) Experimental timeline schematic. (B) Representative *Fos* (red) and *Brs3* (white) expression in the PBN of naïve animals without heat stimulation. (C) Representative *Fos* (red) and *Brs3* (white) expression in the PBN of naïve animals with heat stimulation. (D) Representative *Fos* (red) and *Brs3* (white) expression in the PBN of CFA animals without heat stimulation. (E) Representative *Fos* (red) and *Brs3* (white) expression in the PBN of CFA animals with heat stimulation. (F) Percent of cells expressing both *Brs3* and *Fos* in the left and right PBN (Mixed-effects analysis, effect of hemisphere p=0.3373, effect of injury p<0.0001, effect of stimulation p=0.0366, n=4-5 animals, average of 3-5 sections per animal). (G) Percent of PBN cells co-expressing *Fos* and *Brs3* in the PBN (left and right hemispheres combined) (One-way ANOVA, p=0.0002, with Tukey’s multiple comparisons). All data includes both male and female animals.

To determine whether inflammatory pain similarly alters the functional activity of *Brs3*-expressing PBN neurons, we next performed *in vivo* fiber photometry during hindpaw stimulation before and after CFA administration (**Figure 2A-C**). At baseline, *Brs3* neurons exhibited little to no calcium responses to innocuous light touch (**Figure 2D**), firm static mechanical stimulation (**Figure 2H**), or noxious heat (**Figure 2L**). Three days after CFA injection, calcium responses to innocuous light touch remain unchanged (**Figure 2E-G**). In contrast, *Brs3*-expressing neurons displayed significantly enhanced calcium transients in response to both firm mechanical stiulation (**Figure 2I-K**) and noxious heat (**Figure 2M-O**). Together, these results complement the molecular activation data (**Figure 1**) and demonstrate that inflammation selectively sensitizes *Brs3*-expressing PBN neurons to noxious, but not innocuous, sensory stimuli.

**Figure 2:**
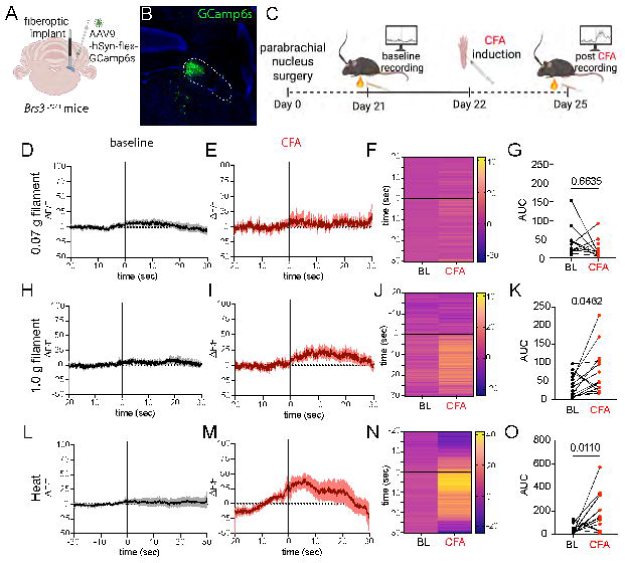
Calcium activity of *Brs3* PBN neurons before and after CFA. (A) Schematic of viral and fiberoptic implant surgery in the PBN. (B) Representative viral expression of GCamp6 in *Brs3* neurons within the PBN (region demarcated by white dashed line). (C) Timeline schematic for the fiber photometry experiments. (D) Averaged calcium transients in response to 0.07 g von Frey filament at baseline and (E) after CFA. (F) Heat map showing averaged responses across animals at baseline and after CFA in response to a 0.07 g filament. (G) Area under the curve summary data for the five seconds directly following stimulation with a 0.07 g filament (paired t-test, p=0.6635, n=14). (H) Averaged calcium transients in response to 1.0 g von Frey filament at baseline and (I) after CFA. (J) Heat map showing averaged responses across animals at baseline and after CFA in response to a 1.0 g filament. (K) Area under the curve summary data for the five seconds directly following stimulation with a 1.0 g filament (paired t-test, p=0.0462, n=14). (L) Averaged calcium transients in response to radiant heat at baseline and (M) after CFA. (N) Heat map showing averaged responses across animals at baseline and after CFA in response to radiant heat. (O) Area under the curve summary data for the five seconds directly following stimulation with radiant heat (paired t-test, p=0.0110, n=14). Solid black line indicates time of stimulus application. Solid trace represents average response, shaded region represents SEM. Each data point in AUC graphs represent responses from a single animal. All data includes both male and female animals.

To evaluate whether *Brs3*-expressing PBN neurons contribute to inflammation-induced behavioral hypersensitivity, we used Cre-dependent inhibitory chemogenetics in CFA-treated *Brs3^Cre^*^/+^ mice (**Figure 3A**). CFA produced robust mechanical allodynia in mice expressing hM4Di; however, transient inhibition of PBN *Brs3*-expressing neurons with the DREADD agonist C21 did not alter mechanical withdrawal thresholds (**Figure 3B**). In contrast, C21-mediated inhibition of *Brs3* PBN neurons significantly increased withdrawal latencies to radiant heat in the Hargreaves assay, reversing CFA-induced heat hypersensitivity (**Figure 3C**). C21 had no effect on mechanical or thermal sensitivity in *Brs3^+^*^/+^ control mice. These findings suggest that *Brs3*-expressing PBN neurons selectively contribute to heat, but not mechanical, hypersensitivity following inflammatory injury.

**Figure 3:**
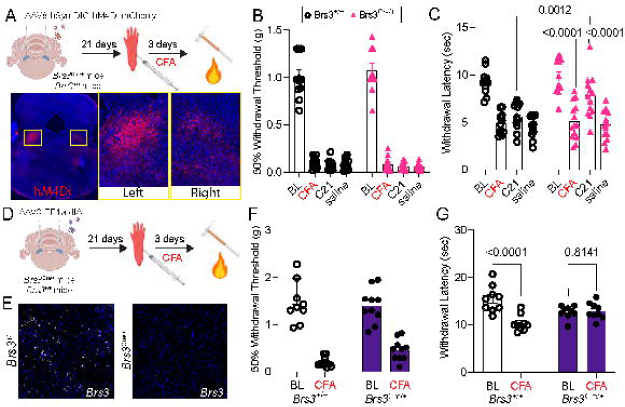
***Brs3*-expressing PBN neurons are necessary for heat but not mechanical hypersensitivity.** (A) Schematic for infecting *Brs3* PBN neurons with Cre-dependent inhibitory chemogenetic (hM4Di) virus. (B) Mechanical withdrawal thresholds before and after CFA in response to intraperitoneal injection of saline or C21 (Two-way ANOVA, effect of time, p=0.2378, effect of genotype, p=0.2199, effect of interaction p=0.8340, n=11-14). (C) Withdrawal latency to radiant heat before and after inhibition of PBN *Brs3* neurons in hM4Di animals treated with CFA (Two-way ANOVA, effect of time p<0.0001, effect of genotype, p=0.0824, effect of interaction, p=0.0147, Tukey’s multiple comparisons, n=11-14). (D) Schematic for genetic ablation of *Brs3*-expressing neurons in the PBN. (E) dTA-mediated ablation of *Brs3* in the PBN. (F) Mechanical withdrawal thresholds before and after CFA in *Brs3^Cre/+^* and *Brs3^+/+^* mice (Two-way ANOVA, effect of treatment p<0.0001, effect of genotype p=0.0996, effect of interaction p=0.1474, n=9). (G) Withdrawal latency on a 52°C hotplate before and after CFA in animals with intact PBN *Brs3* and animals with ablated PBN *Brs3* (Two-way ANOVA, effect of interaction p=0.0021, Uncorrected Fisher’s LSD, n=9). All data includes both male and female animals.

We next tested the hypothesis that permanent genetic deletion of *Brs3*-expressing PBN neurons would prevent the development of CFA-induced pain-like behaviors by using Cre-dependent diphtheria toxin A (DTA) injected bilaterally into the PBN of *Brs3^Cre/+^*and *Brs3^+/+^* mice three weeks before behavioral testing (**Figure 3D**). Efficient genetic deletion of PBN *Brs3*-expressing neurons (**Figure 3E**) had no effect on baseline sensory responses or the development of CFA-induced mechanical allodynia (**Figure 3E-3F**). In contrast, *Brs3*-ablated mice failed to develop heat hypersensitivity after CFA, whereas wildtype controls exhibited the expected reduction in withdrawal latency (**Figure 3F**). Together, these findings demonstrate that *Brs3*-expressing PBN neurons are required for the development and expression of inflammatory heat hypersensitivity while being dispensable for mechanical allodynia.

We next asked whether this apparent heat specificity extended beyond inflammatory pain. Using *in vivo* fiber photometry, we recorded calcium activity in *Brs3*-expressing PBN neurons during hindpaw stimulation before and after spinal nerve ligation (SNL), a model of neuropathic pain (**Figure 4A-B**). Similar to the CFA model, SNL did not alter calcium responses to light or firm mechanical stimulation (**Figure 4D-K**). However, responses to noxious heat were significantly enhanced following SNL (**Figure 4L-O**), indicating that *Brs3*-expressing PBN neurons become selectively sensitized to heat across distinct models of persistent pain.

**Figure 4:**
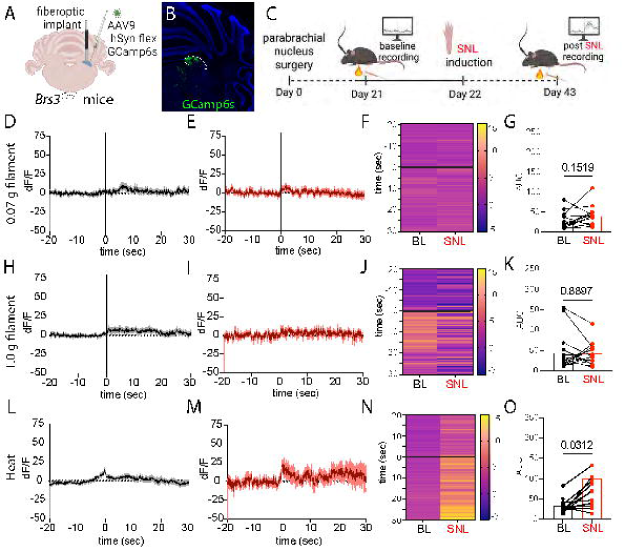
Calcium activity of *Brs3* neurons before and after SNL. (A) Schematic of viral and fiberoptic implant surgery in the PBN. (B) Representative viral expression of GCamp6 in *Brs3* neurons within the PBN (region demarcated by white dashed line). (C) Timeline schematic for the fiber photometry experiments. (D) Averaged calcium transients in response to 0.07 g von Frey filament at baseline and (E) after SNL. (F) Heat map showing averaged responses across animals at baseline and after SNL in response to a 0.07 g filament. (G) Area under the curve summary data for the five seconds directly following stimulation with a 0.07 g filament (paired t-test, p=0.1519, n=14). (H) Averaged calcium transients in response to 1.0 g von Frey filament at baseline and (I) after SNL. (J) Heat map showing averaged responses across animals at baseline and after SNL in response to a 1.0 g filament. (K) Area under the curve summary data for the five seconds directly following stimulation with a 1.0 g filament (paired t-test, p=0.8897, n=14). (L) Averaged calcium transients in response to radiant heat at baseline and (M) after SNL. (N) Heat map showing averaged responses across animals at baseline and after SNL in response to radiant heat. (O) Area under the curve summary data for the five seconds directly following stimulation with radiant heat (paired t-test, p=0.0312, n=14). Solid black line indicates time of stimulus application. Solid trace represents average response, shaded region represents SEM. Each data point in AUC graphs represent responses from a single animal. All data includes both male and female animals.

Finally, we investigated whether the *Brs3*-expressing PBN neurons similarly regulate neuropathic pain-like behaviors using Cre-dependent inhibitory chemogenetics in the SNL model (**Figure 5A-B**). SNL produced robust static mechanical (**Figure 5C**), dynamic mechanical (**Figure 5D**), cold (**Figure 5E**), and heat (**Figure 5F**) hypersensitivity in both *Brs3^Cre^*^/+^ and *Brs3^+^*^/+^ mice. Consistent with our findings in CFA-treated mice (**Figure 3**), C21-mediated inhibition of parabrachial *Brs3*-expressing neurons selectively reduced heat hypersensitivity without affecting static mechanical, dynamic mechanical, cold hypersensitivity, or noxious pin prick (**Figure 5C-F**). Together, these findings identify *Brs3*-expressing neurons as a modality-selective parabrachial population that preferentially regulates heat hypersensitivity across both inflammatory and neuropathic pain states.

**Figure 5:**
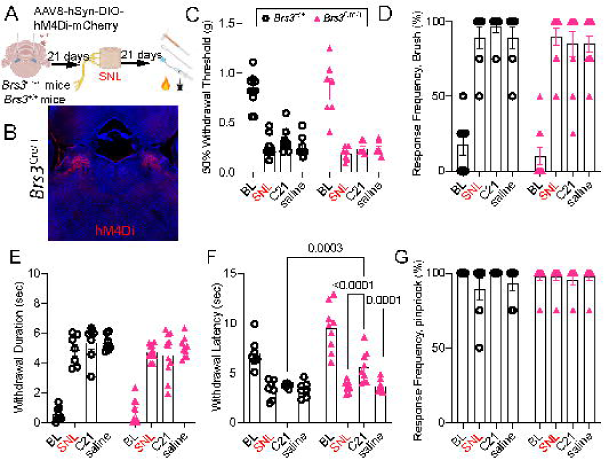
Inhibition of parabrachial *Brs3* selectively alleviates heat hyperalgesia in SNL mice. (A) Schematic of experimental timeline. (B) Representative viral expression of hM4Di in *Brs3* neurons in the bilateral PBN. (C) Mechanical withdrawal thresholds before and after SNL in response to intraperitoneal injection of saline or C21 (Two-way ANOVA, effect of time, p=0.2470, effect of genotype, p=0.0642, effect of interaction p=0.5576, n=7-10). (D) Response frequency to dynamic brush before and after SNL in response to intraperitoneal injection of saline or C21 (Two-way ANOVA, effect of time, p=0.7816, effect of genotype, p=0.5166, effect of interaction, p=0.5099, n=7-10). (E) Withdrawal duration to the acetone droplet before and after SNL in response to intraperitoneal injection of saline or C21 (Two-way ANOVA, effect of time p=0.5343, effect of genotype, p=0.1529, effect of interaction, p=0.4732, n=7-10). (F) Withdrawal latency to radiant heat before and after SNL in response to intraperitoneal injection of saline or C21 (Two-way ANOVA, effect of time, p=0.0010, effect of genotype, p=0.0129, effect of interaction, p=0.0323, Uncorrected Fisher’s LSD, n=7-10). (G) Response frequency to noxious pinprick before and after SNL in response to intraperitoneal injection of saline or C21 (Two-way ANOVA, effect of time, p=0.5361, effect of genotype, p=0.4431, effect of interaction, p=0.1904, n=7-10). All data includes both male and female animals.

## Discussion

The PBN has emerged as a critical hub for pain processing, relaying ascending nociceptive signals to forebrain regions that coordinate the sensory and affective aspects of pain. Recent years have seen a resurgence of interest in the role of the parabrachial nucleus (PBN) in pain processing^22,23,28,30–32,35,36,43,44,49,53–57^. Advancements in technology have enabled rapid experimental progress in dissecting the PBN’s contribution to pain modulation since it was originally identified as a vital node in the pain pathway in the late 1980s^58,59^. New studies have highlighted not only the PBN’s critical involvement in pain but also its remarkable heterogeneity^23,30^. Here, we leveraged recent spatial transcriptomic data^30^ to identify *Brs3*-expressing neurons as a distinct PBN subpopulation that selectively regulates heat hypersensitivity across inflammatory and neuropathic pain states.

Our findings demonstrate for the first time that *Brs3*-expressing neurons in the PBN selectively contribute to heat hypersensitivity during persistent pain. *Brs3*-expressing neurons exhibited increased *Fos* expression following noxious heat stimulation in CFA-treated mice, and enhanced calcium responses to heat following both CFA and SNL, while remaining largely unresponsive under baseline conditions. This selective recruitment suggests that parabrachial *Brs3*-expressing neurons participate in pathological, rather than physiological, processing of thermal nociception. Consistent with this interpretation, inhibiting *Brs3*-expressing neurons reversed heat hypersensitivity without affecting mechanical hypersensitivity in either inflammatory or neuropathic pain models. Together, these findings identify *Brs3*-expressing neurons as a modality-specific PBN population that preferentially regulates thermal hypersensitivity during persistent pain.

Glutamatergic neurons in the PBN are activated by inflammatory and neuropathic pain^23,27,31,32^, but the precise subpopulations contributing to specific types of pain are still being discovered. Glutamatergic tachykinin 1 receptor (*Tacr1*) neurons in the superior lateral PBN are active in response to multiple noxious stimuli, including noxious, but not non-noxious, heat, and inhibition of these neurons reduces inflammatory pain-like behavior ^35^. Parabrachial *Tacr1* neurons are also involved in modulating mechanical and thermal hypersensitivity in the context of neuropathic pain^60^. *Brs3* neurons co-express *Tacr1* and are also located in the superior lateral subnucleus of the PBN (slPBN); therefore, they may represent the noxious heat specific population in this area.

Neuropeptide Y Y1 receptor-expressing parabrachial neurons are also activated by inflammatory and neuropathic pain, and inhibition of these neurons reverses pain-like behaviors in both models^36^. However, while *Npy1r* neurons respond to noxious heat as observed in *in vivo* calcium imaging, modulation of parabrachial *Npy1r* neurons does not alleviate thermal hyperalgesia associated with CFA^36^. It is likely that the *Brs3* subset of *Npy1r* neurons are the ones responding to heat, but they may represent a large enough population to have a modulatory effect when manipulated. We hypothesize that *Brs3*-expressing PBN neurons represent a unique subset of glutamatergic neurons that encapsulate established neural subpopulations already implicated in inflammatory pain - *Tacr1* and *Npy1* - and may serve as a key integrative hub that drives heat hyperalgesia in the context of persistent pain.

Beyond the parabrachial nucleus, *Brs3*-expressing PBN neurons project to the hypothalamus, thalamus, lateral preoptic area, and amygdala, suggesting their potential involvement in integrating nociceptive signals with affective and homeostatic processing^41^. Pain is intricately involved in homeostasis-it reflects an adverse condition that affects autonomic functions and causes a behavioral response^1^. Given that terminal regions *Brs3*-expressing PBN neurons project to also regulate energy metabolism, emotion, and negative affect, it is likely that parabrachial *Brs3* neurons have additional physiological roles beyond pain modulation^41^. Parabrachial projections to different brain regions are involved in discrete aspects of the pain experience^55^. Thus, future studies should investigate whether downstream parabrachial *Brs3* projections contribute to the emotional and aversive aspects of pain as well as autonomic functions affected by pain conditions.

### Limitations and Future Directions

One limitation of this study is the time course of the chosen models. CFA and SNL models were chosen to represent distinct persistent pain conditions with a heat phenotype. CFA pain-like behavior peaks at 3 days post injection, but mechanical and heat allodynia persists for up to two weeks^61^. Allodynia associated with SNL persists longer but eventually recovers^62^. Future studies should include more timepoints, as *Brs3* neurons in the PBN do not seem to influence normal heat sensitivity at baseline, and they may be involved in discrete time periods, like the development or maintenance of different pain conditions. Additionally, to further investigate the importance of parabrachial *Brs3*-expressing neurons to heat allodynia in the context of chronic pain, other persistent pain conditions with temperature related allodynia should be explored.

Additionally, the upstream circuitry that recruits *Brs3*-expressing PBN neurons during persistent pain remains unknown. One possibility is that these neurons receive direct input from modality-selective spinal projection neurons, extending the concept of labeled lines for pain beyond the spinal cord into the brainstem. Consistent with this idea, a population of glutamatergic spinal neurons expressing *ErbB4* was recently shown to selectively mediate heat hypersensitivity^63^. Future studies should determine whether a dedicated circuit exists linking peripheral afferents expressing the transient receptor potential family member V1 (TRPV1) channel ^64,65^ to *ErbB4*-expressing spinal neurons to *Brs3*-expressing PBN neurons.

Because both the PBN and *Brs3*-expressing neurons are involved in various physiological processes, it will be critical to assess potential off-target effects by monitoring vital signs and other readouts of homeostatic behavior (e.g., feeding) during *Brs3* neuron stimulation. Similarly, the contribution of *Brs3* PBN neurons to non-reflexive measures of pain, such as aversion-based behavioral assays, could also clarify whether *Brs3* neurons influence the affective dimensions of pain.

## Conclusion

The PBN is a highly complex hub that processes and relays a vast array of sensory inputs, including pain. There is growing interest in dissecting the contributions of specific excitatory subpopulations within the PBN to distinct aspects of the pain experience. We found parabrachial *Brs3* neurons are preferentially involved in modulating heat hypersensitivity, suggesting that different PBN populations process persistent pain-related mechanical and thermal hypersensitivity separately. These data support the idea that pain is not a monolithic experience but instead consists of multiple parallel pathways.

## Acknowledgements

We thank Dr. Richard Palmiter for generously donating the original *Brs3^Cre^* mice. BioRender.com was used for making schematics. This work was funded by National Institutes of Health awards F32NS128392 (HNA), K00NS124190 (TSN), RF1NS131165 (RK), PhRMA Foundation Postdoctoral Fellowship in Drug Discovery 1335819 (EJRP), and a Development Grant from the American Neuromuscular Foundation (TSN).

## Disclosures

R. Khanna is the co-founder of Regulonix LLC, a company that develops non-opioid drugs for the treatment of chronic pain. No other authors have declared conflicts of interest.

## Author contributions

HNA, TSN, and RK developed the concept. HNA designed and conducted the experiments, completed the data analysis, and wrote the manuscript. TSN helped conduct *in situ* experiments and EJRP conducted SNL surgery and behavior. NAD and NKG assisted with tissue processing and conducted confirmatory immunohistochemistry and RNAscope. All authors had the opportunity to discuss the results and comment on the paper.

## Notes

### Summary of Updates

New data, including a new pain model being added

